# Development of a label-free microfluidic impedimetric immunoassay for anti-SPAG16 antibody

**DOI:** 10.1101/2021.02.25.432859

**Authors:** T Froyen, F Vreys, L Slaets, V Somers, R Thoelen

## Abstract

A recently found biomarker regarding multiple sclerosis, namely the anti-SPAG16 antibody (Ab), could be a potential way for early detection, prognosis and diagnosis of said autoimmune disease. Merging electrochemical analysis with a microfluidic system is a novel approach, which avoids the use of labelling steps as seen in traditional ELISA immunoassays. In this study, aluminium interdigitated electrodes on polystyrene-coated PET foils were implemented in a microfluidic flow cell to bind and detect SPAG16 Abs by impedimetric measurements. The coated PET foils showed a clear affinity for the fusion protein SPAG16-THIO and thioredoxin (THIO). Determining sensitivity and specificity of antibody-antigen binding using a microfluidic ELISA immunoassay has revealed the test to be unreliable by showing no linear pattern of a dilution series of the standard and producing skewed inconsistent results. The impedimetric analysis showed opposite results of what one would expect. The systems efficiency is in need to be revised and optimised before undergoing actual diagnostic tests. Further advancement could be done by reducing leakage, securing a more stable entrance for injection and circumventing the occurence of air bubbles in the wells.

**SAMENVATTING:** Multiple sclerosis (MS) is een auto-immune ziekte, gekenmerkt door inflammatie van het centraal zenuwstelsel en demyelinisatie van axonen. Een recent gevonden proteïne, namelijk het anti-SPAG16 antilichaam, is bewezen een biomerker te zijn voor MS. Met behulp van een elektrochemisch analytisch systeem gecombineerd met microfluidica zou men anti-SPAG16 antilichamen kunnen opsporen voor vroegtijdige detectie, prognose en diagnose. Het grote voordeel tegenover traditionele ELISA testen is het elimineren van *labelling* stappen waardoor de immunoassay goedkoper en gebruiksvriendelijk wordt. In deze studie werden overlappende aluminium elektrodes op polystyreen-gecoate PET-plaatjes geïmplementeerd in een microfluidische flow cell voor binding en detectie van anti-SPAG16 antilichamen door middel van impedimetrische metingen. De gecoate PET-plaatjes vertonen een duidelijke affiniteit voor 0,1 μg/ml SPAG16 en THIO. Het bepalen van sensitiviteit en specificiteit van antilichaam-antigen binding, gebruikmakende van een ELISA immunoassays, gaf aan dat de test onbetrouwbaar is doordat geen duidelijk patroon voor standaardoplossingen zichtbaar was en resultaten vertekend en inconsistent bleken te zijn. De impedimetrische analyse vertoonde tevens onbetrouwbare resultaten, waarbij een omgekeerd effect werd geobserveerd van wat er in theorie zou moeten gebeuren. De efficiëntie van het systeem moet herzien en geoptimaliseerd worden voordat men het voor diagnostische testen kan gebruiken. Verdere vorderingen zouden gerealiseerd kunnen worden door lekkages te reduceren, steviger bevestigen van de PDMS-ingang en de ontwikkeling van luchtbellen in de wells te vermijden.

## INTRODUCTION

Immunoassays are reliable tools to identify target molecules and their concentration in a given sample. The accuracy of these techniques is guaranteed because of the high selectivity and specificity of antigen – antibody interactions. Impedance-based sensors could trump conventional immunoassays and be further improved by implementing microfluidic channels. The development of bioanalytical systems aims to improve the cost, ease-of-usage, reproducibility and reliability without being dependent on the need for any specialized laboratory. Improvement of these characteristics will ensure a better high-throughput capacity and therefore faster identification of diseases (1).

Enzyme-linked immunosorbent assays (ELISAs) have become widely used diagnostic bioassay designs for proteins, peptides, hormones and other biomarkers since the technique was first published in 1971 (2). ELISA has four different main formats; direct, indirect, sandwich, and competitive, which differ in the number and order of reagents used. Each of these forms contains an enzyme-conjugated antibody which will induce a quantifiable change in the environment proportional to the antigen-concentration (3). Direct (enzyme-conjugated primary antibody), indirect (enzyme-conjugated secondary antibody) and sandwich ELISA have a progressive linear relationship between concentration and target-binding, meaning higher concentrations of the target will result in a higher magnitude of a predetermined signal (4, 5). However competitive assays are set up in a way in which higher concentrations will result in lower levels of signals. Labelled add-in antigens compete with the non-labelled sample antigens resulting in a decreased signal with increasing concentrations of the sample antigen. The regression of the concentration-signal curve will therefore be reverse linear opposed to its other three forms (6). These conventional ELISA techniques require several reagents, each has to be incubated for X amount of time, making it time-consuming procedures. Depending on the form of ELISA, these reagents include sample antigens, labelled antigens, primary/secondary enzyme-conjugated antibodies, blocker, stopping solution, etc. Apart from the number of reagents, there are other limitations of ELISA that should be eliminated to create the best results for diagnostic purposes. Incorrect modification of labelled add-in antigens, as seen in competitive ELISAs, results in aberrant outcomes (7). Spectrophotometric ELISAs can experience some interference caused by a deficient light-source density (turbid colouring) and therefore limiting the accuracy and sensitivity (8). Although the ELISA has been a helpful tool in ubiquitous clinical laboratories, the modern rate of diagnostic tests requires a less labour-intensive, complicated and bulky alternative with lower costs (9).

A possible and promising future for the next generation immunoassays contains electrochemical sensors which not only tackle the limitations of conventional ELISAs but are also less expensive, labour-intensive and time-consuming. This paper will focus on impedance-based measurements, which is a method that uses detection via electrical impedance tomography. The characteristics of the electrode-electrolyte interface and the analytes in the sample will influence the impedance between the electrodes in a microfluidic cell. Because of the deviation of impedance occurring during target-binding on the electrode, the change can accurately determine concentrations of a given sample using a calibration curve of said sample (9). This technique is considered a label-free immunoassay by eliminating the use of enzyme-conjugated antibodies, and the reagents following these antibodies, making it easier to implement it in research projects and diagnostics than conventional enzyme-immunoassays. Furthermore, by making the impedance-associated immunoassay a microfluidic device, it will gain a few additional advantages in comparison to traditional immunoassays. Sample and reagent fluids are able to flow in specially engineered channels ranging in several microliters, allowing more economical usage of healthcare resources. Miniaturization of the procedure reduces volumes of samples and reagents, risk of contamination and enhances detection limits, making the effects synergistic with those of the impedance-based immunoassay (10). Combining microfluidics with electrochemical immunoassays seems to have a promising role in the future of laboratory research since it enhances user-friendliness and lowers cost-rates (11). In this work, platforms of polyethylene terephthalate (PET) or glass both containing gold (Au) or aluminium (Al) microelectrodes coated with polystyrene act as a base for target-binding and measurements. Modification of the electrode surface with polystyrene creates an environment known to immobilize antigens. Polystyrene is a biologically inert thermoplastic consisting of aromatic rings connected to hydrocarbon chains that immobilize proteins and peptides using covalent bonds. Polystyrene provides multiple anchoring points for amino acids found on proteins/peptides ensuring entrapment (12). Aspecific binding locations are blocked with non-fat milk (NFM) to prevent false-positive results before the eventual antibody is added to the mixture (13). These processes occur in a microfluidic device, approximately 5 mm in diameter and 2,1 mm high, where samples and reagents flow through the device to provide the electrodes with the previously mentioned components. The frequency-dependent resistance to the current flow in between the electrodes is measured and interpreted to obtain our final results.

The investigated analyte in our set-up, namely anti-SPAG16 Abs, is a recently found biomarker present in cerebrospinal fluid (CSF) and plasma samples of patients with multiple sclerosis (MS). MS is a chronic inflammatory disease prominent in the CNS causing loss of myelin sheaths and neurodegeneration. The autoimmune nature of MS is responsible for these inflammatory effects and the onset of further disease progression (14). De Bock et al. hypothesized that astrocytes upregulate SPAG16 expression as a consequence of proinflammatory mediators as seen in MS sites (15). The expression of SPAG16 is not restricted to the central nervous system, moreover it is mostly expressed in the testis owing to the fact that it plays an important role in ciliary and flagellar motility of germ cells (16, 17). Monoclonal antibodies can be directed against multiple similar epitopes found in different isoforms of the protein. These epitopes are apparent in a reactive region of SPAG16 Abs consisting of 121 amino acids which can be proven by using a basic local alignment tool (Supplementary Fig. S1) (15, 18).

## MATERIALS AND METHODS

### Microfluidic system ELISA

4 × 1.7 × 0.01 cm PET foils were coated with 0.25% polystyrene (solvent: 99.8% toluene, Sigma-Aldrich). A multi-layered setup was designed for ELISA, intended for holding 50 μl volume of fluids. A more spacious design allows better detection of colour change opposed to smaller channels, therefore we layered multiple materials to create a complex consisting an incoming channel, a cylindrical container and outflowing channel. The polystyrene-coated PET foil is connected, via double-sided tape, to a bulkier PMMA plate. A second double-sided tape holds on to a non-coated pet foil, which is further covered with a 4 × 1.7 × 0.01 cm double-sided Kapton tape meant to stabilize a polydimethylsiloxane (PDMS) block with a hole overlapping with each circular end of the Kapton tape as an entry/exit point. The different layers were cut by a Speedy 100R Laser cutter (Trotec). Plastic tubes connect the whole on one end with the reagents, samples and buffers and on the other end with a High Precision Multichannel Dispenser (ISMATEC) at 5% flow rate. All can be viewed in Fig. 1A.

**Fig. 1:**
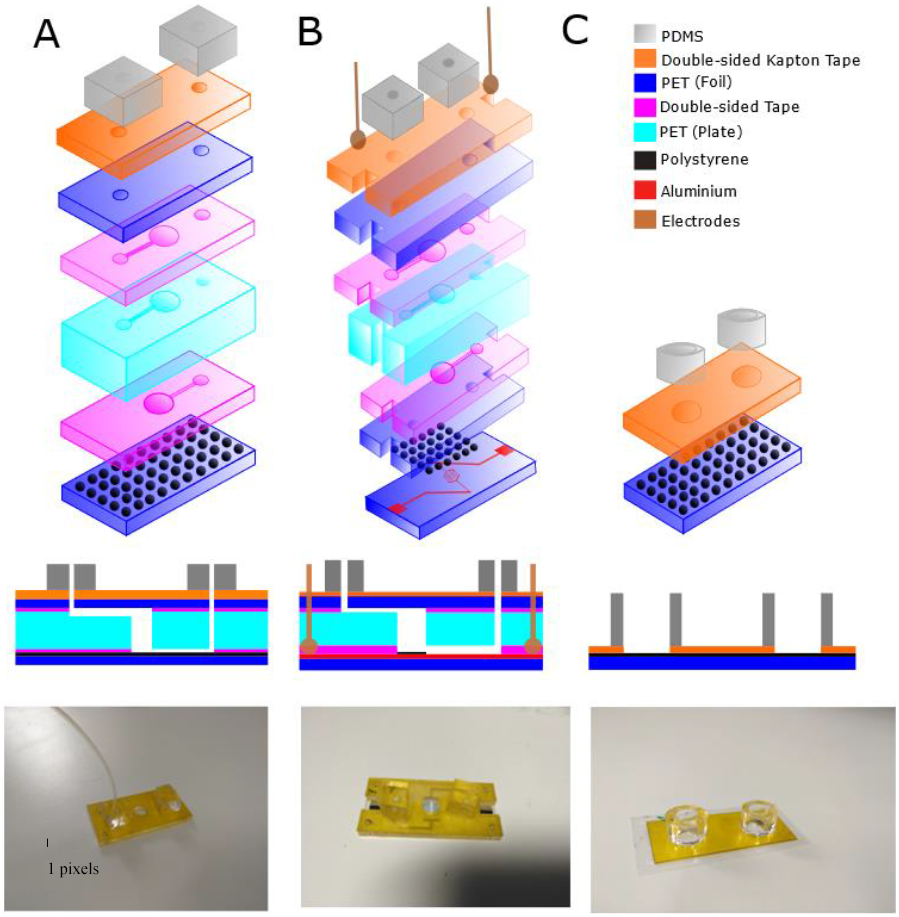
Microfluidic setup for ELISA-, impedance- and contact angle measurements. Microfluidic systems visualised by a 3D- and 2D scheme followed by an additional photo of the product for ELISA (A), impedance spectrometry (B) and contact angle measurements (C). Legend displays the different layers in function of colours.

### Microfluidic system impedance spectroscopy

The same system as ELISA was used, except for the aluminium-grafted modification of the PET foil, acting as an interdigitated electric circuit that can be connected with electrodes (Fig. 1B).

### Microfluidic system contact angle measurements

Polystyrene-coated PET foils were directly covered with double-sided Kapton tape with two holes as a mean to expose the polystyrene-coated surface. A well was created from each hole using a cylindrical block of PDMS able to hold at least 50 μl of fluid. The general principle of the design can be seen in Fig. 1C.

### ELISA

Multiple polystyrene-coated PET plates were incubated overnight at 4°C with 1μg/ml and 0.1 μg/ml protein (SPAG) in coatingsbuffer (9.6pH) (stock concentration: 1.2 mg/ml SPAG16 and 965.6 μg/ml THIO) and 1 μg/ml and 0.1 μg/ml THIO as a control. Washing buffers consisted of 1xPBS and 0.1% PBS-Tween (PBST). Aspecific binding sites of the polystyrene layer were blocked using 2% non-fat milk in PBS (MPBS) for 1h at 37°C before adding 20 μl plasma samples (1:100 in 2% MPBS) for 1h at room temperature. Binding of monoclonal Abs was detected using rabbit HRP-conjugated anti-human IgG (1:2000 in % MPBS) for 1h at room temperature. Substrate 3,3’,5,5’-tetramethylbenzidine (TMB) was used as initiator of colour development. 2M H_2_SO_4_ was used after 10 minutes to stop the reaction, which was then transferred in a sterile 96 well plate (Greiner Bio-One) and read at 450 nm with an ELISA plate reader (Biomed, Diepenbeek).

### Impedance Tomography

Much like ELISA, polystyrene-coated PET plates were incubated overnight at 4°C with 1 μg/ml protein and 0.1 μg/ml (SPAG) in coatingsbuffer (9.6pH) (stock concentration: 1.2 mg/ml SPAG16 and THIO 965,6 μg/ml). Washing buffers consisted also of 1xPBS and 0.1% PBS-Tween (PBST). Aspecific binding sites were blocked using 2% MPBS for 1h at 37°C before adding 2 μl plasma samples (1:100 in 2% MPBS) for 1h at room temperature. Impedance was measured after the well was filled with PBS at 20°C using frequencies between 0.1-20 kHz using an Iviumstat electrochemical interface (Ivium Technologies). The method regarding impedance tomography is visualised in Fig. 2.

**Fig. 2:**
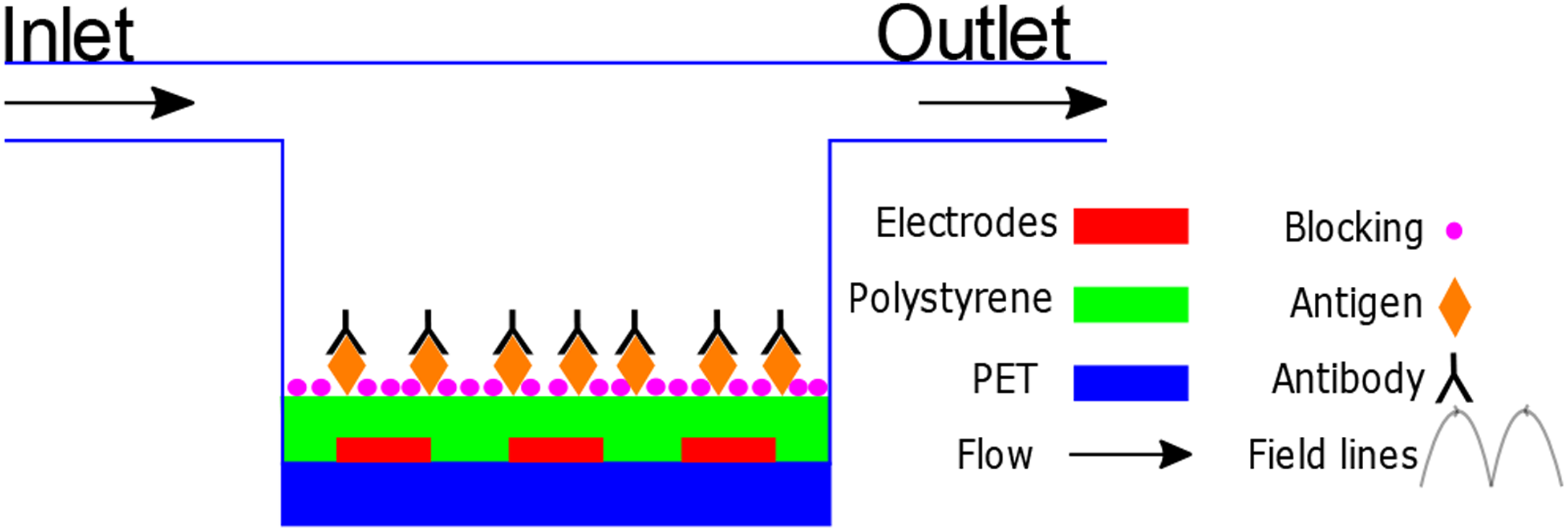
Principle of impedance-based measurements in a label-free immunoassay: Interdigitated electrodes placed on polystyrene-coated PET foils manifest field lines when a current runs through them. The impedance of this system is correlated with the amount of bound antigen-antibody complexes on the surface, which can be translated in the exact concentration of said antigen.

### Contact Angle Measurements

The contact angle of SPAG16 (1*10^−1^, 1, 10 and 100 μg/ml), THIO (1*10^−1^, 1, 10 and 100 μg/ml), coatingsbuffer, marvel and the bare polystyrene surface on PET foils was measured using Optical Contact Measuring and Contour Analysis Systems (OCA) 15 Plus. Each reagent was spotted two times on multiple polystyrene-coated PET plates and incubated 16h at 5°C.

### Statistical analysis

Statistical analysis was performed using JMP Pro 14 software (SAS). Analysis of variance was used to compare different concentrations of SPAG16 during contact angle measurements. P-values ≤ 0.05 were considered statistically significant (* p ≤ 0.05; ** p ≤ 0.01; *** p ≤ 0.001). Data are mean ± SD

## RESULTS

### Contact Angle Measurements

To validate proper immobilisation of anti-SPAG16 Abs on the polystyrene-coated PET surface, the contact angle of MiliQ water droplets was measured to determine a noticeable change in surface properties (19). The angle of the droplet can be used as a measure of hydrophobic or hydrophilic tendencies of surfaces and could additionally be used to estimate the degree of coating needed to ensure sufficient anti-SPAG16 Ab-surface binding. The bare polystyrene surface exhibits a poor wettability, which establishes the hydrophobic character of polystyrene-coated PET and sets a baseline for the further experiments. A positive control, namely coatings buffer, was used to prove the ability to change the wettability of the surface. The surface becomes hydrophilic in contrary to its non-modified version, indicating a change of surface traits.

Binding of blocker buffer on the substrate is necessary to avoid skewed data and false-positive results, therefore verifying the affinity of non-fat milk to the surface will assure the exclusion of non-specific binding to the substrate. The surface, where MPBS was spotted, showed a significant difference in wettability, indicating reliable blocking of patches currently not occupied by antibodies.

A range of concentrations (1*10^−1^, 1, 10 and 100 μg/ml) of SPAG16 antigen and thioredoxin (THIO) were incubated on polystyrene-coated PET surfaces to determine their degree of surface modification compared to bare polystyrene. All measured concentrations of SPAG16 antigen and THIO exhibit increased levels of hydrophilicity, which exaggerates with rising concentrations. Significant differences of surface wettability were found between 0.1 μg/ml and 10 μg/ml, as well as 0.1 μg/ml and 100 μg/ml SPAG16 antigen. These findings indicate lesser binding occurs using lower amounts of antibodies, like 0.1 μg/ml. The same results can be seen with a concentration of 1 μg/ml compared to 10 and 100 μg/ml SPAG16 antigen (Fig. 3). However, the two lower concentrations still give an adequate change in surface wettability to bind a sufficient amount of antigen to perform our test with. No significant difference in wettability was found between 0.1 μg/ml and 1 μg/ml, meaning the lowest tested concentration could be used for further experiments.

**Fig. 3:**
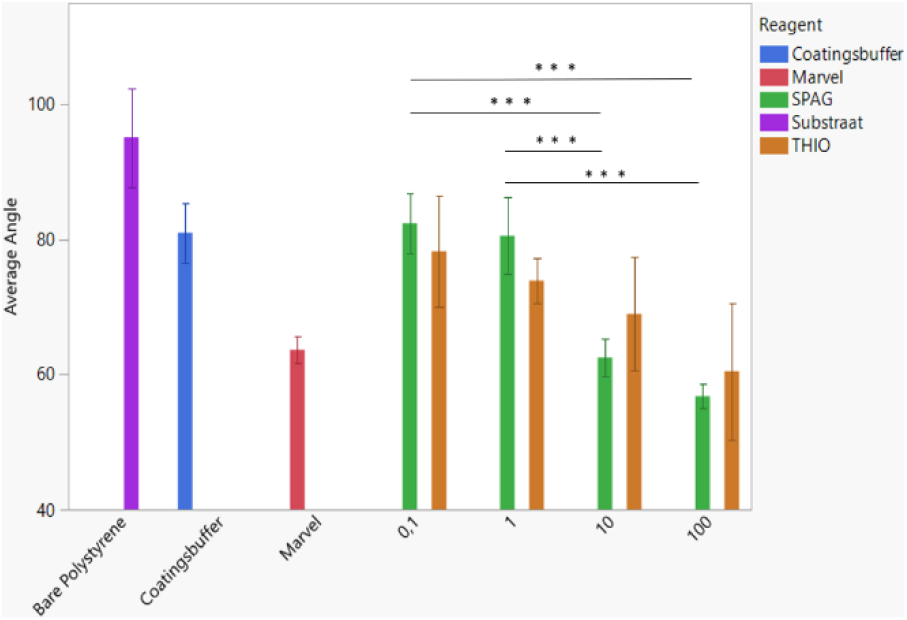
Surface contact angle affected by different coatings on polystyrene-coated PET foils: Droplet contact angle measurements of bare polystyrene-coated surface (n=12) with or without additional Marvel (n=6), coatingsbuffer (n=8), SPAG16-(0.1 μg/ml: n=8, 1 μg/ml: n=10, 10 μg/ml: n=9, 100 μg/ml: n=10) and THIO antigen (0.1 μg/ml: n=8, 1 μg/ml: n=8, 10 μg/ml: n=10, 100 μg/ml: n=10) coating after overnight incubation. Data are mean ± SD. ***P<0.001, one-way analysis of variance.

THIO follows a similar pattern as SPAG16 antigens. A more hydrophilic character is observed, that becomes more apparent with higher concentrations. Likewise with anti-SPAG16 Abs, the preferable change occurs when the foil is incubated with 10 μg/ml or 100 μg/ml. Loo ing at the lower concentrations (0.1 and 1 μg/ml) used in our experiments, we see a greater change in wettability of THIO than SPAG16, meaning THIO has a stronger affinity to the polystyrene-coated surface. The difference between the mean angle between both antigens is respectively 4.16° for 0.1 μg/ml and 6.65° for 1 μg/ml. However, the results involving THIO show a greater rate of variability making the angles less predictable, which translates to a non-uniform coating of the polystyrene-coated PET foils. The same range of variability could explain the lack of significance found between the different concentrations, apart from 0.1 μg/ml and 100 μg/ml along with 1 μg/ml and 100 μg/ml THIO. The increase in magnitude of hydrophilicity with larger concentrations is visualised in Fig. 4. A vast difference in contact angles of the two highest and two lowest concentrations is observed, assisting the results of our previous seen graph. An increase of surface-bound antigens flattens out the droplet, while less binding of antigens augments the degree of bulging indicating a more hydrophobic character as seen in a bare polystyrene surface.

**Fig. 4:**
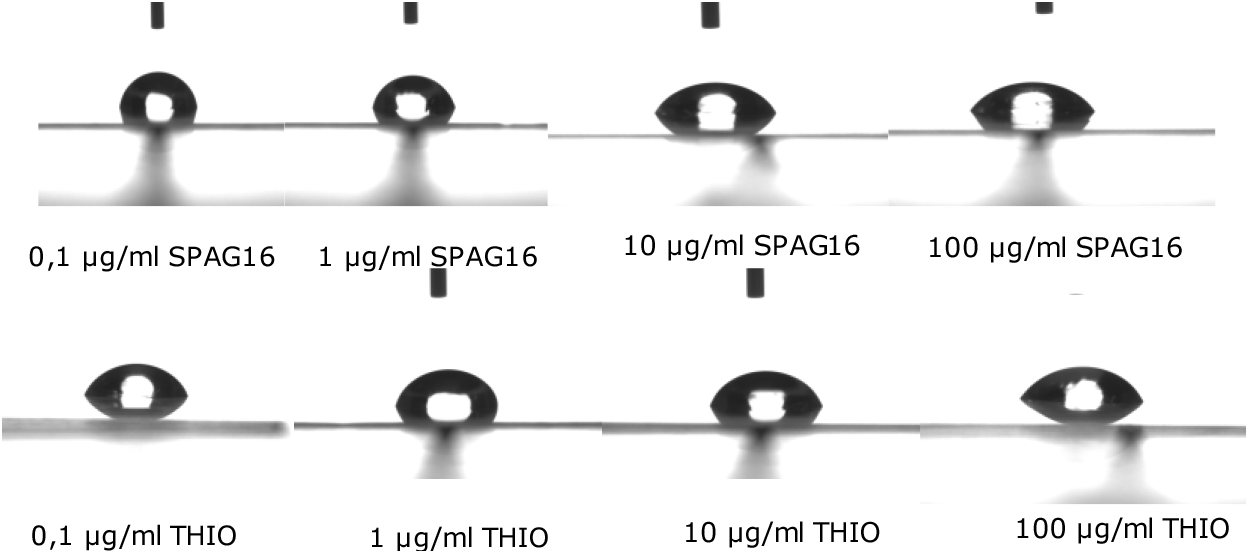
Visualisation of changes in material wettability due to different degrees of surface modification: Photos of water droplets on polystyrene-coated surface modified with SPAG16- and THIO antigen in different concentrations, which changes the curvature of the droplet. Higher concentrations antigen flatten out the droplet, indicating more bound antigen to the surface and therefore a higher degree of modification.

**Fig. 5:**
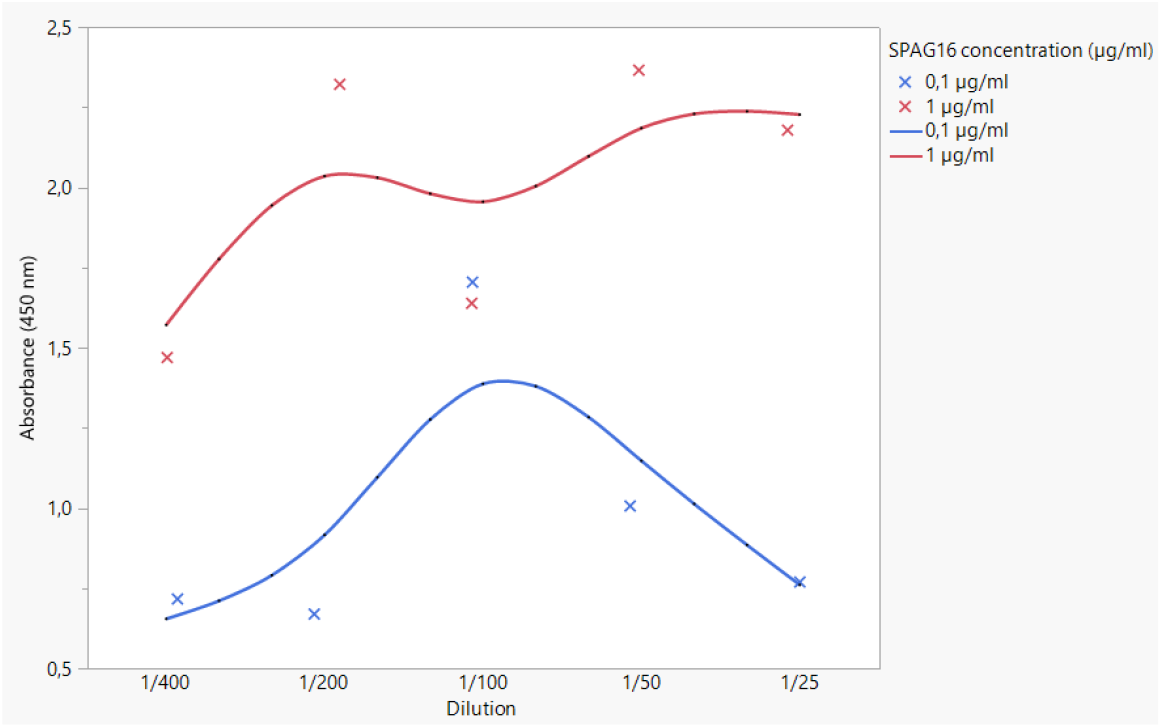
Absorbance of different SPAG16 concentrations in standard solutions to determine sensitivity: ELISA (enzyme-linked immunosorbent assay) measurements using absorbance at 450 nm on different dilutions (1/25, 1/50, 1/100, 1/200 and 1/400) of 0.1 μg/ml (n=5) and 1 μg/ml (n=5) SPAG16.

**Fig. 6:**
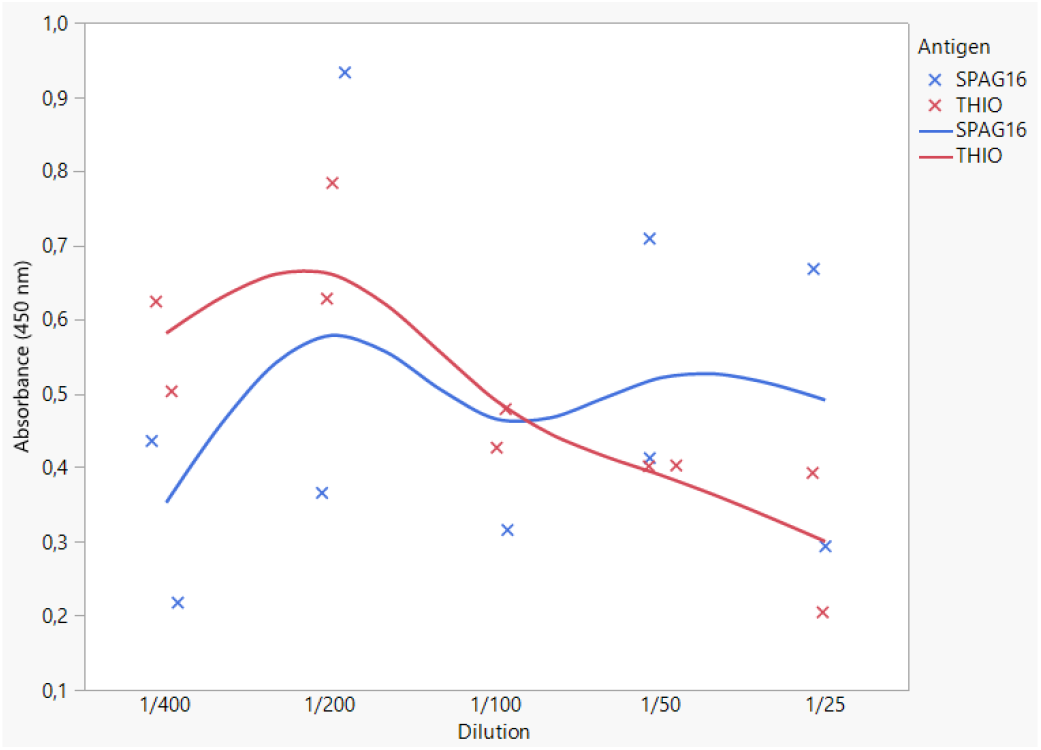
Absorbance of SPAG16 and THIO standard solutions to determine specificity: ELISA (enzyme-linked immunosorbent assay) measurements using absorbance at 450 nm on different dilutions (1/25, 1/50, 1/100, 1/200 and 1/400) of 1 μg/ml SPAG (n=9) and THIO (n=10).

### ELISA

To establish the sensitivity of our microfluidic ELISA setup to detect SPAG16 antigen, two different concentrations were used which were previously proven with contact angle measurements to bind on the polystyrene-coated surface. Concentrations 0.1 μg/ml and 1 μg/ml SPAG16 antigen were thus used in standard solutions with dilution factors of 1/25, 1/50, 1/100, 1/200 and 1/400 to determine if both concentrations could be detected.

Larger concentrations have more potential for colour development and exhibit as consequence more absorbance of light. Therefore, a normal ELISA standard curve has a positive linear or slightly exponential course. However, in this paper the axiom doesn’t seem to uphold itself because both curves diverge strongly from the expected pattern. In theory, the less diluted sample should display a larger absorbance, however the concentration of 0.1 μg/ml SPAG16 antigen starts to decline after peaking at dilution factor 1/100. Hence, this gaussian curvature shows the same rate of absorbance between dilution 1/400 up till 1/200 and dilution 1/50 up to 1/25, making the determination of sample concentrations by interpolation seem ambiguous. As expected, the concentration of 1 μg/ml SPAG16 antigen has an overall higher absorbance than 0.1 μg/ml antigen with the biggest difference of absorbance seen in dilution 1/200 by 1.625. The curve itself has a more positive course as the standard solution becomes less diluted, nevertheless the graph is not ideal because of the waving pattern. The dilution factor 1/100 makes a downfall in absorbance, passing dilution 1/200 in the lower end of the curve. Apart from these remarks, both concentrations could be detected by ELISA, thus making a first estimate of the test’s sensitivity.

Specificity was determined by comparing colour development of 1 μg/ml SPAG16 with 1 μg/ml THIO. Contact angle measurements showed the affinity of THIO to polystyrene-coated PET foils making the test more susceptible to false positives. The presence of both antigens were thus measured using anti-SPAG16 antibodies.

These results show also a divergent pattern opposed to the predicted values as seen in traditional ELISA standard curves. The absorbance of antigen SPAG16 diminishes suddenly after dilution 1/200 following with a waving pattern as if dilution of our concentration doesn’t seem to affect the colour development of the solution. Just like SPAG16, THIO reaches its peak at 1/200 before the following values drop drastically with decreasing dilution, which is the exact opposite of what we would expect. The lower concentrations THIO seem to exhibit more absorbance than SPAG16 by a difference of 0.2365 in absorbance at dilution 1/400. This gap closes up at dilution 1/200 with a difference of 0.0565 and reaches an equilibrium between the two antigens around dilution 1/100. After crossing paths, SPAG16 antigens gain a greater absorbance than THIO between dilution 1/100 and 1/25 opposed to the smaller concentrations. Concentration 1 μg/ml SPAG16 antigen diluted 1/50 shows more absorbance than THIO by a difference of 0.1585 which escalates to 0.182 at dilution 1/25.

### Impedance Spectrometry

To obtain a label-free immunoassay, impedance is used to determine the concentration of SPAG16. The setup is tested by using 1 μg/ml SPAG16 standard solutions diluted by factor 1/25, 1/50, 1/100, 1/200 and 1/400. The largest difference between the impedimetric results was found in between frequencies 100-598 Hz (Fig. 7), therefore we analysed the following impedance measurements using a frequency of 245 Hz.

**Fig. 7:**
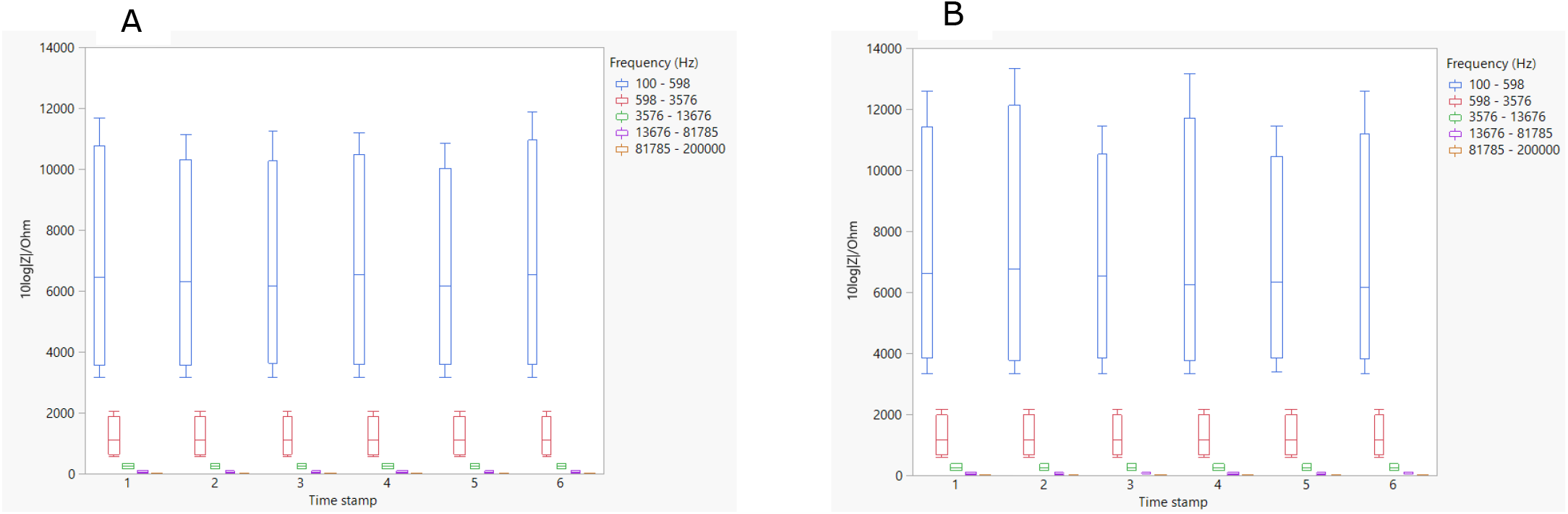
Deviations in impedance regarding different intervals of frequencies using SPAG16- and THIO antigen coating. Impedance tomography measurements at frequencies 0.1-20 kHz for (A) SPAG16 antigen (n=18 for each time stamp) and (B) THIO antigen (n=18 for each time stamp).

The overall pattern of SPAG16 is relatively the same between different dilutions, however the bars of the histogram decreases in impedance after every step of the protocol (Fig. 8). Our blank consists of the microfluidic flow cell filled with PBS buffer as a starting reference. The expected outcome would be an continuous increase in impedance after binding of antigen, blocking buffer and finally the antibodies. However, the blank in each of our tests is the highest value on the plot, while binding of antibodies in the last step, seems to have the smallest degree of impedance. The difference in resistance is the highest between the blank and after binding of our antigen. The impedance-based sensor shows minor decreases of resistance in the last three steps relatively to the blank. The average registered impedance measurement difference between the blocker step and binding of the antibody for dilution 1/25 is 172,80 Ω with and standard deviation of 120 Ω starting at 5106,88 Ω of the blocker step. Dilution 1/50 exhibits between the last two steps an average of 250.26 Ω with a standard deviation of 137.91 Ω starting at 6613.11 Ω, dilution 1/100 has an average of 113.47 Ω with a standard deviation of 72.69 Ω starting at 5741.24 Ω, dilution 1/200 has an average of 285,97 Ω with a standard deviation of 134 Ω starting at 5740.29 Ω and the last dilution impedimetric measurement averages at 209.66 Ω with a standard deviation of 152.24 Ω starting at 5171.42 Ω, thereby giving not much margin of error for the higher dilutions to make a clear distinction between concentrations of actual clinical samples.

**Fig. 8:**
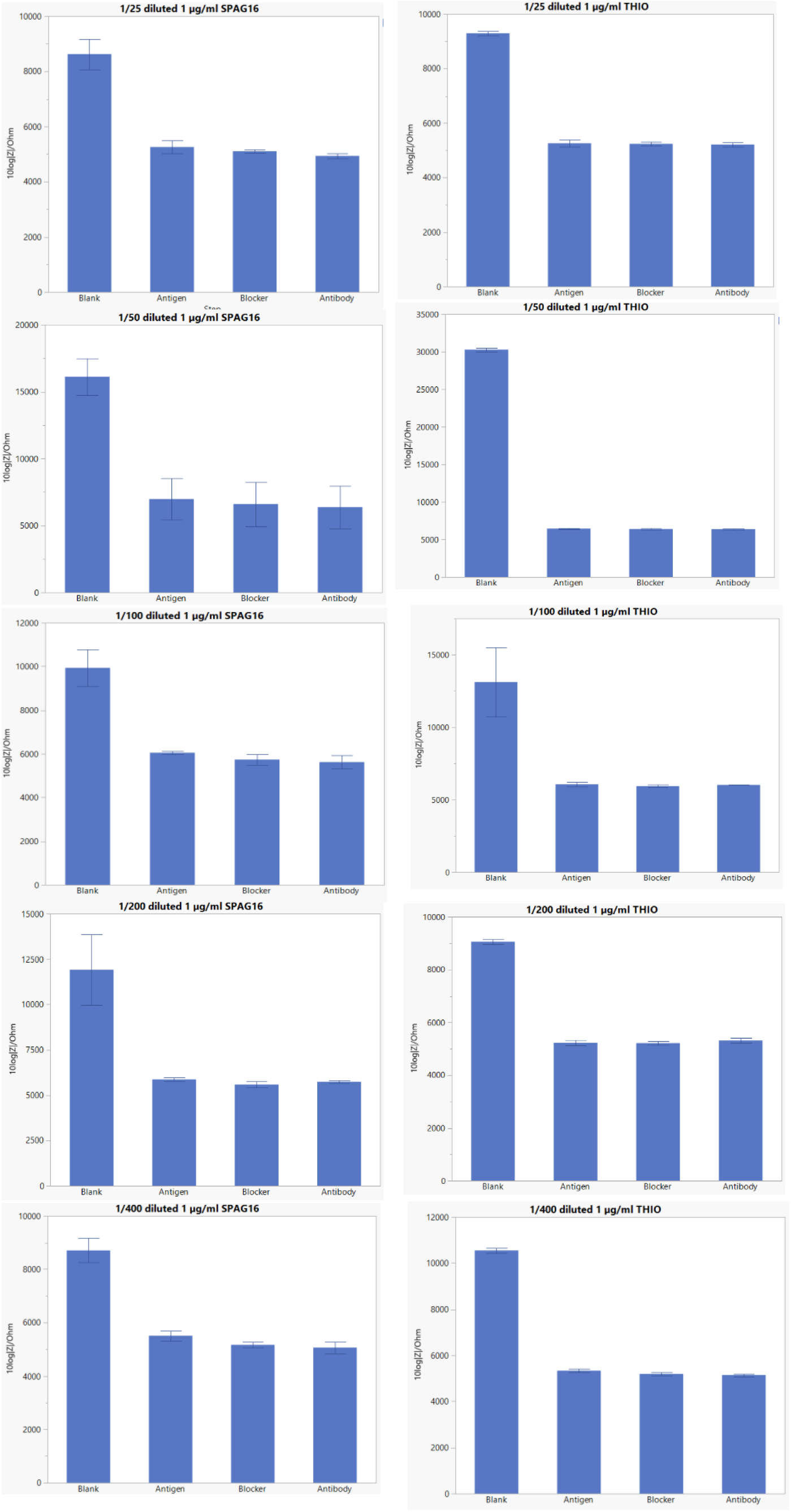
Impedance measurements of SPAG16- and THIO after major steps in the protocol using different dilutions. Impedance tomography was used to measure dilutions 1/25, 1/50, 1/100, 1/200 and 1/400 before and after the addition of SPAG16- or THIO antigens (incubation overnight, THIO: n=5 every step per dilution, SPAG n=10 every step per dilution), blocker buffer (1h incubation) and antibodies (1h incubation). Data are mean ± SD.

The control THIO has a similar pattern in all different dilutions as SPAG16, although the impedance measurements, regarding the last three steps of the protocol, seem to be stagnant in contrary to the decrease as seen with SPAG16 antigen (Fig. 8). Measurements show that the blank has the highest impedance with a rapid downfall after coating the surface with THIO antigen. The blocking- and antibody-binding step fall into the same range of impedance as SPAG16 with an average of 5599.44 Ω for the blocker step and an average of 5616.12 Ω for the antibody-binding step, hinting at a non-optimal specificity regarding antibody-antigen binding.

## DISCUSSION

To make an impedimetric microfluidic immunoassay to work, all parts must function correctly to minimise interference of the outside of the flow cell and ensure tight sealing to prevent any seepage of fluids. Binding of the reagents must be as effectively as possible to heighten the level of impedance between the electrodes of the electrical circuit. In theory, the more antibody-bound antigens on the polystyrene-coated surface should increase the resistance of current flow, which is detectable if the system possesses enough sensitivity (20). The results of our experiments have shown an opposite effect by a decrease in impedance by every step. The system should be revised by confirmation of the biological effectiveness and capability for detection by the use of a traditional non-microfluidic ELISA to ensure the fault lies within the mechanics of our impedimetric microfluidic system instead of flawed execution of the actual protocol. After verification of correct operating, improvements concerning the construction of the microfluidic cell is advised because of adhesive failure of the different layers. The biggest problem was securing the PDMS blocks on the Kapton tape during our experiments, because loss of adherence between these two layers caused major leakages, prevention of flow and filling of the well. Other causes of leakages could be designated to faulty stacking of the different layers, creating smaller passageways and openings with diminished flow as a result. Another problem was air bubbles arising in our well preventing binding of antigens, blocker buffer, antibodies and interference of impedance measurements. The source of air remaining in our well is most likely due to suboptimal passage of fluids. Faltering movements of the fluid is a good indication of possible air bubble formation in the microfluidic well and should be dealt with before going any further with the protocol.

The same problems can be attributed to the skewed results found in the traditional ELISA test to determine sensitivity and specificity. Additionally, non-uniform sticking of PDMS blocks to the exterior of the flow cell could create little pockets in which fluids could be left behind and potentially generate false-positive results if not washed away properly. Further, retrieval of volumes for detection in a plate reader was inconsistent and affected the initial concentrations in the wells making the measurements unreliable. This again, can be attributed to faltering of the fluids which in turn makes their velocity unpredictable and hard to manage. The low specificity could therefore be caused by the little pockets in PDMS and aberrant patterns of standard solutions by improper passage.

The polystyrene-coated PET foil has proven to be an effective modification to bind proteins to the surface. The blocking buffer, SPAG16- and THIO all bind to the exterior in a uniform way, meaning errors caused by improper binding of proteins to the surface can be ruled out as a responsible factor for skewed measurements. Moreover 0.1 μg/ml and 1 μg/ml SPAG16 show to have the same degree of affinity to the surface, indicating a promising potential for sensitivity especially when backed up by detection of all dilutions of 0.1 μg/ml SPAG16 during the ELISA test. However, we still need to take the unreliable character of the latter test into account to make such assumptions. No contact angle measurements were done of polystyrene-coated PET surfaces with implemented aluminium circuits to validate an absence of interference to bind proteins by the metal aluminium. To ensure the same reliability of protein-binding on aluminium-grafted polystyrene-coated surfaces as well as bare polystyrene surfaces, further tests can be done in the future to obtain a more well-rounded view of the impedimetric microfluid immunoassays capability.

All the previously mentioned problems involving the microfluidic aspect of the immunoassay should be addressed, revised and resolved before continuing the pursuit of a label-free assay since unreliable results caused by microfluidic errors automatically make the impedimetric measurements obsolete. Correct stacking of layers should receive special attention to avoid improper passage, while PDMS blocks should be firmly secured to the entrance of the flow cell by a better adhesive or clamps. Leakages and formation of air bubbles are the biggest flaws of the impedimetric test but are resolvable, thereby ensuring a potential future of our label-free immunoassay. Improvements of principle itself could be done by changing the use of a direct ELISA test to a sandwich ELISA test that uses aptamers instead of immunoglobulins (21). This way, we’ll create a label- and immobilisation-free assay further reducing the needed time-frame significantly due to its ability to be long-termed stored, while its regeneration capabilities could further enhance the cost-effectiveness (22). This could be considered after dealing with the current issues.

## CONCLUSION

The results proved the impedimetric microfluidic immunoassay, described in this study, to be untrustworthy and unpredictable. The construction of the microfluidic flow cell seems to be the biggest culprit at the moment, causing leakages and bubble formation. The overall system still has a long way to go and is in need for several adjustments but that doesn’t ta e away its potential to replace the traditional ELISA test. The future operating version will have a remarkable beneficial influence in clinical laboratories, not only for SPAG16 but other antigens as well.

## Acknowledgements

The author is grateful to Dr. Laura de Bock (Biomed, Diepenbeek) to provide SPAG16- and THIO antigen samples along with the necessary material for the ELISA immunoassays.

## Author contributions

FV, LS, VS and RT conceived and designed the research. TF and FV performed experiments and data collection. FV and LS aided with analytical systems. FV developed the analytical tool. Statistical analysis and writing of the paper were performed by TF.

## SUPPORTING INFORMATION

**Fig. S1:**
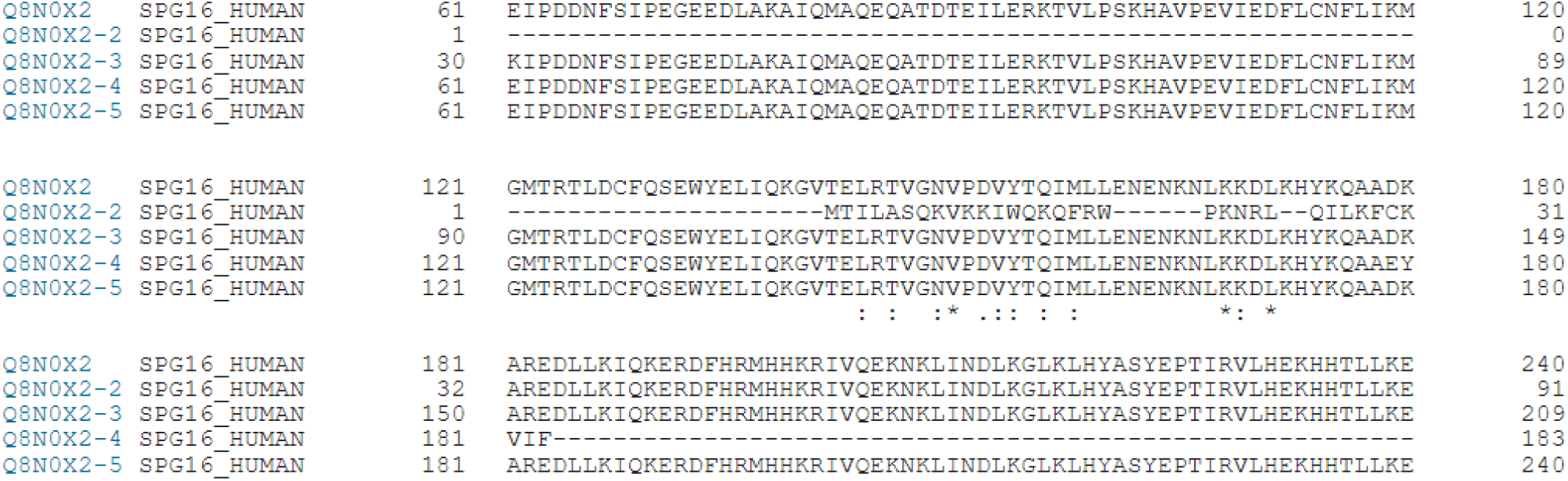
Similarities between amino acid sequences of various SPAG16 isoforms. Similar amino acid sequences between isoforms of SPAG16 act as a reactive region consisting of multiple epitopes for SPAG16 antibody binding.

